# A GABAergic Maf-expressing interneuron subset regulates the speed of locomotion in *Drosophila*

**DOI:** 10.1101/421057

**Authors:** H Babski, C Surel, S Yoshikawa, J Valmier, J.B Thomas, P Carroll, A Garcès

**Affiliations:** Inserm U1051, Institute for Neurosciences of Montpellier, University of Montpellier, 34295 Montpellier, France; Molecular Neurobiology Laboratory, The Salk Institute for Biological Studies, La Jolla, CA 92037, United States.

**Keywords:** Central Pattern Generators, intersectional genetics, locomotion, Maf Transcription factor, interneurons, MNB progeny neurons, Transcription factor combinatorial code, *Drosophila* larva.

## Abstract

Interneurons (INs) coordinate motoneurons activity to generate adequate patterns of muscle contractions, providing animals with the ability to adjust their body posture and to move over a range of speeds. In the *Drosophila* larvae several IN subtypes have been morphologically described and their function well documented. However, the general lack of molecular characterization of those INs prevents the identification of evolutionary counterparts in other model animals, limiting our understanding of widespread principles ruling neuronal circuits organization and functioning. Here we characterize a highly restricted neuronal subset expressing the Maf transcription factor Traffic Jam (TJ). We found that TJ^+^ neurons are highly diverse and their activation using intersectional genetics disrupted larval body posture and locomotion speed. We also showed that a small subset of TJ^+^ GABAergic INs, singled out by the unique expression of *Per*, *Fkh*, *Grain* and *Hlh3b*, a molecular signature reminiscent to V_2b_ INs in vertebrate, impacted the larvae crawling speed.

## Introduction

The wiring and functioning of the neuronal circuits that provide animals with the ability to move over a range of speeds have been extensively studied (*1*). In mammals as in invertebrates the speed of locomotion is regulated by central pattern generator (CPG) neuronal circuits which coordinate, via motoneurons (MNs), the sequential activation of muscles (*2-5*). The rhythmic bursts of MN activity are controlled by local excitatory and inhibitory interneurons (INs) located in the ventral part of the mammals spinal cord and in the invertebrates Ventral Nerve Cord (VNC). These INs can be categorized in sub-classes on the basis of their connectivity patterns, physiological properties such as neurotransmitter used, and by the set of specific transcription factors (TFs) they express. The diversity of INs is best exemplified by a recent study showing that V_1_ IN class in mammals can be fractionated into 50 distinct subsets on the basis of the expression of 19 TFs (*6*). The combinatorial expression of TFs by subsets of INs thus provides a powerful and systematic tool for assessing genetically, when transgenic lines are available, the function of highly restricted subpopulations of INs. This assessment can be done using intersectional genetics, through the controlled expression of ion channel proteins that regulate INs activity (*7, 8*). Such an approach used in mouse and in *Drosophila* has proved instrumental in dissecting the core logic of the CPG circuits that generate the rhythm and pattern of motor output (*3, 5, 9*). Still, in light of the remarkable diversity of INs in the vertebrate spinal cord (*6, 10*) and in the *Drosophila* nerve cord (*11, 12*), a thorough description of the functioning of the CPG regulating locomotion in animals is far from complete.

A class of segmentally arrayed local premotor inhibitory INs named PMSIs (for period-positive median segmental INs) was recently found to control the speed of larval locomotion by limiting, via inhibition, the duration of MN outputs. The PMSIs have been proposed to be the fly equivalent of the V_1_ INs in the mouse. Thus PMSIs in *Drosophila* and V_1_ INs in vertebrates may represent a phylogenetically conserved IN population that shape motor outputs during locomotion (*5*). In the years following the characterization of the PMSIs, specific IN subtypes contributing to the diversity of locomotor behaviours in the *Drosophila* larvae have been identified, providing a wealth of information on each IN subpopulation, their function, morphology and connections established (*5, 13-15*). However, little is known about the combinatorial expression of TFs within these different IN subtypes identified so far and this lack of knowledge impedes cross species comparisons thus limiting our understanding of the common principles of CPG organization in vertebrates and invertebrates.

Here we investigate in the *Drosophila* larva the role of a small pool of highly diverse INs (23/hemisegment) expressing the evolutionary conserved TF Traffic Jam (TJ), the orthologue of MafA, MafB, c-Maf and NRL in the mouse. Interestingly, MafA, MafB and c-Maf are expressed by restricted subpopulations of ventral premotor INs in the developing mouse spinal cord (*6, 10, 16*) but the function of these INs subtypes is to date unknown. To characterize TJ-expressing INs we generated a *TJ-Flippase* line and used intersectional genetics to activate TJ^+^ subpopulations depending on their neurotransmitter properties. We found that manipulation of these IN subsets modulates larval locomotor behaviours. Our results also showed that activation of a restricted subpopulation of GABAergic/*Per*^+^/TJ^+^ (3 INs/segment), belonging to the PMSIs group of INs and known as MNB progeny neurons, significantly impacts the crawling speed of the larvae.

## Results

### Traffic Jam is expressed in a restricted subpopulation of neurons located in the VNC and involved in larval locomotion

We initiated a detailed analysis of the transcription factor (TF) Traffic Jam (TJ) expression in the embryonic and larval nervous systems, using a previously characterized TJ-specific antibody (*17*) and an enhancer trap for *TJ* (*TJ-Gal4*) (*18*). During embryogenesis, TJ expression is first detected in late stage 12 (st 12) in few cells in the brain and in 12 to 15 cells/hemisegment in the ventral nerve cord (VNC) (Supp. Fig.1A). Co-immunostaining with the glial marker Repo showed that TJ is exclusively expressed in neurons and excluded from glia cells (Supp. Fig.1B). Using *TJ-Gal4::UAS-GFP* we found that *TJ-Gal4* faithfully recapitulates TJ expression in all embryonic (Supp. Fig.1C) and larval stages analysed (Fig.1C-G). Closer analysis of TJ expression over time showed that TJ is consistently found in a subset of 29 neurons per hemisegment in the VNC abdominal region (A2-A6) from st17 to L3 larval stages (Fig.1A, A’, B, B’) and excluded from sensory neurons (data no shown). We used *TJ-Gal4::UAS-H2AGFP* in combination with anti-TJ immunostainings to establish a precise topographic map of TJ^+^ neurons in second instar larvae, a stage representative of the continuous expression pattern of TJ (Fig.1 C-G, C1-G1).

**Fig. 1:**
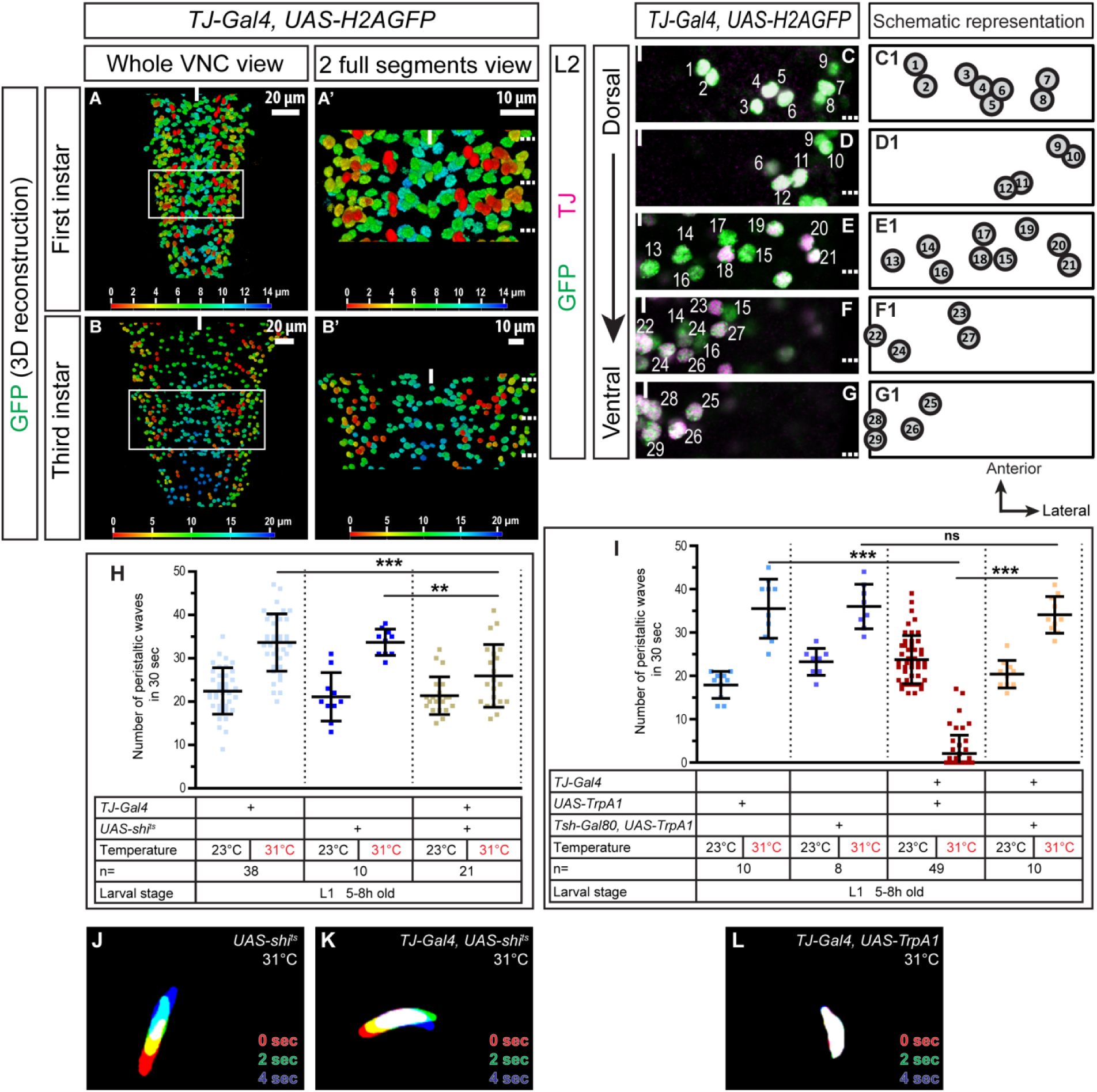
TJ^+^ neuronal population is required for proper larval crawling. **A, A’, B, B’** 3D reconstruction of whole VNC (**A, B**) and 2 full segments (**A’, B’**) of first (**A, A’**) and third (**B, B’**) instar larvae expressing nuclear GFP under the control *TJ-Gal4*. Colour scale is the z-axis scale; most dorsal cells are red, most ventral are blue. White scale is the x/y-axis scale. **C-G** Staining of a second instar larva VNC for TJ (magenta) and *TJ-Gal4* expression reported by nuclear H2AGFP (green). Totality of TJ^+^ cells are shown in dorsal (**C**) to ventral (**G**) panels. Dashed lines on the right-hand side of the panels indicate segment boundaries and the full line the midline. A unique hemisegment is shown in each panel. Anterior of the VNC is up. **C1-G1** Schematic representation of one hemisegment showing stereotyped ventral-dorsal and medial-lateral cell position in first and second instar larvae (cell positions may slightly change in third instar). **H-I** Number of peristaltic waves per 30 seconds at permissive (23°C) and restrictive temperature (31°C). Inhibition of the entire TJ^+^ population (second beige bar, **H**) causes a slight decrease in the number of peristaltic waves. Activation of the entire TJ^+^ population (second red bar) causes a drastic decrease in peristaltic waves (**I**). This decrease is no longer visible upon activation of the TJ^+^ neurons in the brain (second salmon pink bar, **I**). For **H** and **I**, each single point represents recording of a single first instar larva. Error bars indicate the SD and n the number of larvae tested. Statistical analysis: One way ANOVA. ***p≤0.001, **p≤0.01, ns=not significantly different. **J-L** Superimposition of three consecutive time frames (0, 2 and 4 seconds) showing the postures of 5h old larvae: control larvae (**J**), upon inhibition of the TJ^+^ neuronal population (**K**) and upon activation of the TJ^+^ population (**L**).

Next, we decided to explore the function of TJ^+^ neurons in larval locomotion using the *TJ-Gal4* driver as a tool to either inactivate or activate the entire TJ^+^ neuronal population. We inactivated neurons by expressing thermosensitive shibire (*shi*^*ts*^) (*19*); neuronal activation was achieved by expressing *TrpA1* (*20*). Inactivation of the entire TJ^+^ population led to a slight decrease in the number of larval peristaltic waves (Fig.1H – second beige bar). This decrease seemed caused by a general disorganization of the peristaltic waves, with the segments of the larvae failing to contract in the coordinated, sequential way (Fig.1J and 1K, video 1). Activation of the TJ^+^ neurons had more drastic effects, with almost complete abolition of locomotion (Fig.1I – second red bar) and larvae displaying a complete paralysis which we named “spastic paralysis”. This phenotype was characterized by immobility, tonic contraction of all body segments and a drastic shortening of the whole larval body length (Fig.1L, video 2). When placed back at permissive temperature (23°C), the larvae resumed normal locomotion, proving that this spastic paralysis phenotype is fully reversible. TJ is expressed in a restricted number of neurons in the cerebellar lobes. To assess the role of these neurons in the spastic paralysis phenotype, we restricted the expression of *TrpA1* to the TJ^+^ cerebellar lobes neurons using *Tsh-Gal80* (*21*) in combination with *TJ-Gal4* and *UAS-TrpA1*. Under these conditions locomotion appeared normal (Fig.1I – second salmon pink bar), arguing that TJ^+^ neurons in the cerebellar lobes do not have a function in locomotion in *Drosophila* larva.

We thus conclude that TJ^+^ neurons within the VNC are part of a neuronal circuit controlling *Drosophila* larval crawling and that normal function of TJ^+^ neurons is to maintain proper muscle contraction and peristaltic wave propagation during locomotion.

### Activation of TJ^+^ neurons in the VNC using an intersectional genetic approach leads to spastic paralysis of the larvae

To further characterize the identity of TJ^+^ neurons regulating crawling behaviour in the larva we developed an intersectional-based genetic approach, using candidate *LexA* drivers to express a *LexAop-FRT-stop-FRT-dTrpA1* transgene (generous gift from Y. Aso, Janelia Research Campus) in combination with a source of Flippase. To specifically target TJ^+^ neurons we generated a *TJ-Flippase* line. Briefly, within a ∼40kb backbone fosmid carrying all the endogenous regulatory elements of *tj*, we substituted the full open reading frame and 3’ UTR sequences of *tj* with an optimized *Flippase* (*FlpO*) by in *vivo* homologous recombination in *Escherichia coli* (*22*) (see Materials and Methods for more details). We first characterized in “flip-out” experiments the accuracy and efficiency of *TJ-Flp* using *Act>Stop>Gal4::UAS-CD8-GFP* and found from embryonic stage 15 onward that only TJ^+^ cells are GFP-labelled (supplementary Fig.2A). We quantified the efficiency of “flip-out” events in TJ^+^ neurons and found from hemisegment to hemisegment and within all specimen analyzed (n=4) that more than half the TJ^+^ neurons (64%) are already recombined in L1 larva (Fig.2A-C) and 89% in young L2 stage (Fig.2D-F). These numbers demonstrate the accurate recombination triggered by *TJ-Flp*. An examination of developing egg chambers of the ovary, a structure in which TJ has been reported to be specifically expressed by somatic cells (*17*), revealed that all TJ^+^ follicular and border cells are GFP-labelled thus confirming the accuracy of *TJ-Flp* (supplementary Fig.2B-C). Finally, we used *Tsh-LexA* to express a *LexAop-FRT-STOP-FRT-dTrpA1* transgene in combination with *TJ-Flp,* anticipating that activation of TJ^+^ neurons in the VNC only should give rise to the drastic spastic paralysis phenotype described above. We found it was indeed the case in all L1 larvae tested (n=12) (Fig.2K, video 3).

**Fig. 2:**
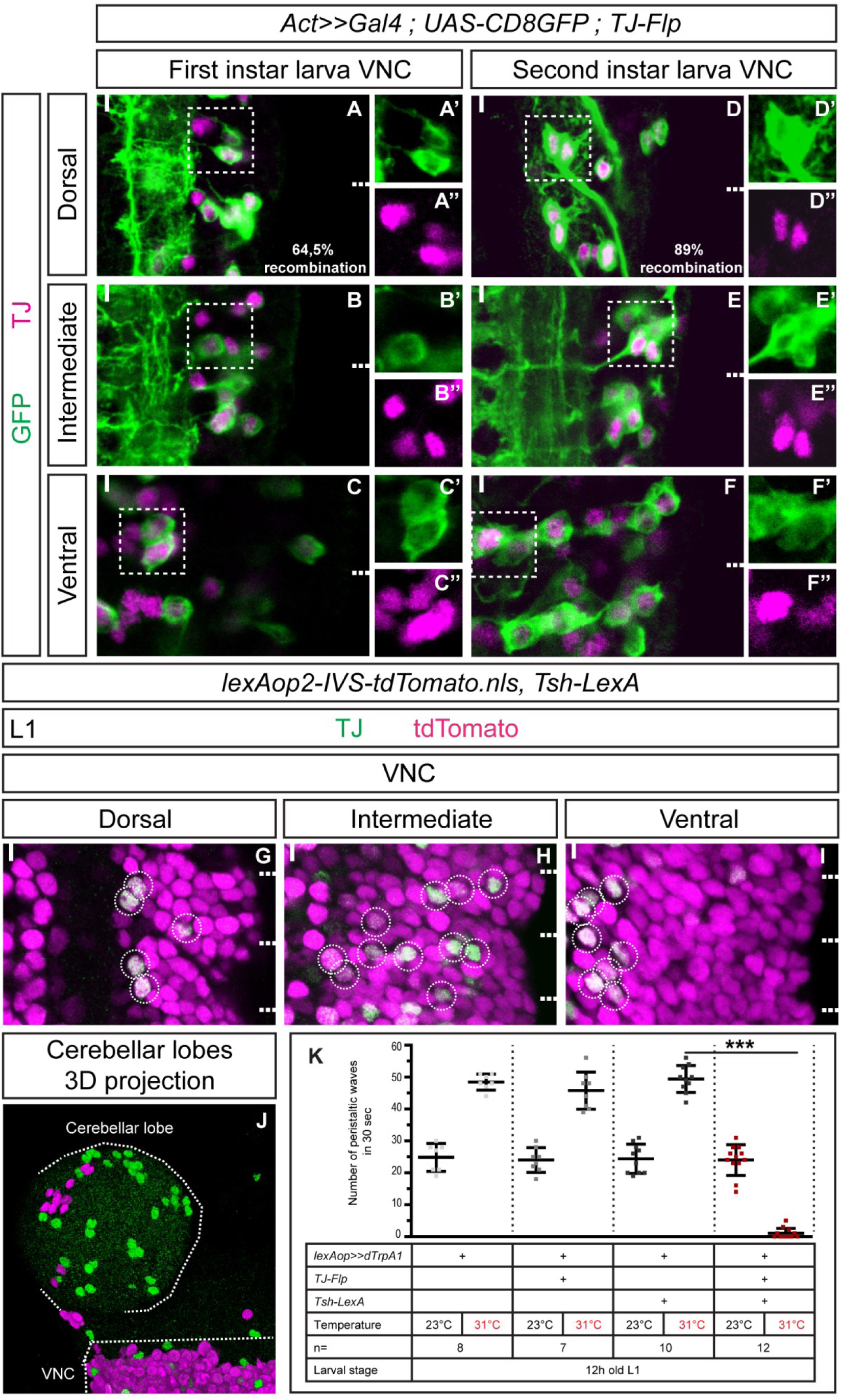
*TJ-Flp* line allows accurate targeting of TJ^+^ neuronal subpopulations. **A-F** Staining for TJ (magenta) and recombined cells (expressing *TJ-Flp* –green) in first (**A to C**) and second (**D to F**) instar larva VNC. Percentages of recombination (calculated as follow: *TJ-Flp*^+^ cells/total TJ^+^ cells*100) are indicated in respective first panels. **G-J** Staining for TJ (green) and *Tsh-LexA* driving an *nls-tdTomato* (magenta) in first instar larva VNC (**G-I**) and in first instar larva cerebellar lobe (**J**). All TJ^+^ cells in the VNC are *Tsh-LexA*^+^ while no co-localisation is found in the brain. **K** Number of peristaltic waves per 30 seconds at permissive (23°C) and restrictive (31°C) temperatures. Upon activation of the TJ^+^ within the VNC only (second red bar), we recapitulate the behaviour observed when the entire TJ^+^ population is activated (presented in Fig.1I). Each single point represents a single 12h-old first instar larva. Error bars indicate the SD and n denotes the number of larvae tested. Statistical analysis: One way ANOVA. ***p<0.001.

Collectively, these results show that the *TJ-Flp* line we generated is an accurate and powerful genetic tool to genetically manipulate TJ^+^ neurons, thus validating our intersectional-based genetic approach.

### TJ^+^ motoneurons activation only partially impacts larval crawling

We next asked whether the spastic paralysis phenotype we observe upon activation of the whole TJ^+^ population would actually be caused by *TJ-Gal4* driving in MNs. Using pMad and Eve as reliable molecular markers for MNs, we found TJ expression from st13 onward in 3 CQ/Us’ MNs namely U1, U2 and U5 (asterisks in Fig.3 A2, A3, B2, B3 and C2, C3) plus another putative MN located in the vicinity of the CQ MNs (full arrowhead in Fig.3 C3). Co-labelling using *Isl-?myc* and pMad also revealed TJ expression in 2 MNs located laterally in the dorsal and intermediate regions of the VNC that are putatively part of the ISNd MNs pool (Fig.3 C2 - inset). We noticed that TJ was neither expressed in aCC nor in any RPs MNs (RP1-5) (Fig.3A1 and C1 – curly brackets) nor in SNa MNs (supplementary Fig.4A1, A2) nor in type II, octopaminergic *Tdc2*^+^ MNs (supplementary Fig.4B).

**Fig. 3:**
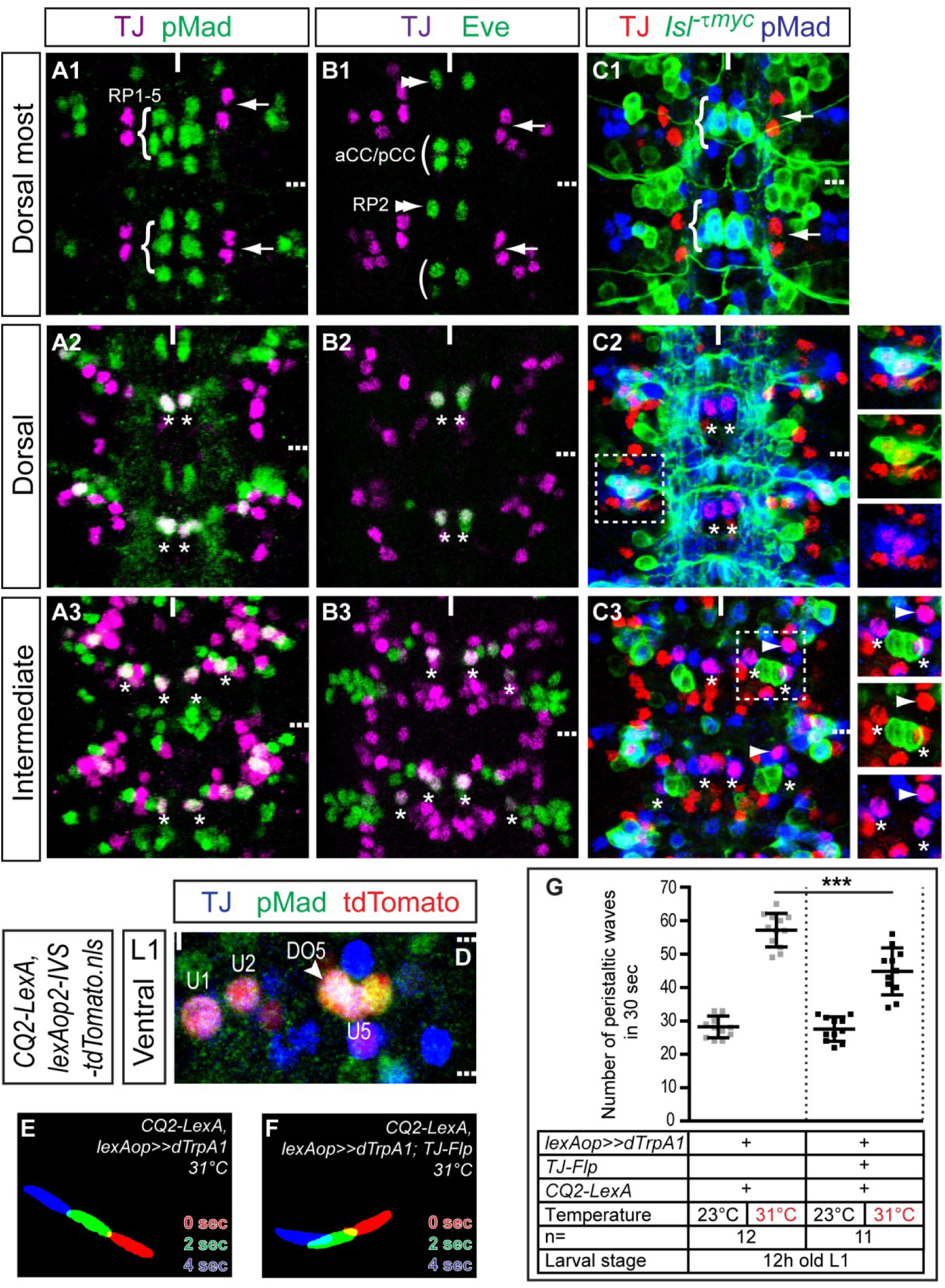
Modulation of TJ^+^ MNs activity moderately impairs larval locomotion. **A1-C3** Representative views of two segments of stage 16 embryo VNC stained for TJ (magenta) and motoneuron marker pMad (green) (**A1-A3**), TJ (magenta) and Eve (green) (**B1-B3**) and TJ (red), pMad (blue) and *Isl*^*-τmyc*^ (green) (**C1-C3**). Cells are shown from dorsal-most (**A1, B1, C1**) to intermediate (**A3, B3, C3**) positions. TJ is excluded from RP1-5 (**A1**, curly brackets) and expressed in U MN (**A2** and **A3**, asterisks). Arrows in **A1** highlight the dorsal-most pair of TJ^+^ neurons. TJ is excluded from aCC/pCC (**B1**, curly brackets) and RP2 (**B1**, double arrowhead) and expressed in U MNs (**B2** and **B3**, asterisks). Arrows in **B1** highlight the dorsal-most pair of TJ^+^ neurons. TJ is expressed in 2 dorsal lateral *Isl*^*+*^ motoneurons (**C2**, inset), in the *Isl*^-^ U MNs (**C2** and **C3**, asterisks) and in 1 TJ^+^ pMad^+^ MN that innervates DO5 (**C3**, full arrowhead). Arrows in **C1** highlight the dorsal-most pair of TJ^+^ neurons and asterisks in **C2** and **C3** show U MNs. **D** Staining for TJ (blue), Eve (green) and endogenous nls-tdTomato driven *CQ2-LexA* in a first instar larva VNC. *CQ2-LexA* drives in TJ^+^ U MNs U1, U2 and U5 as well as the MN innervating DO5 muscle (**D,** full arrowhead) and 1 non-identified TJ^+^ IN (**D**, arrow). **E-F** Superimposition of three consecutive time frames (0, 2 and 4 seconds) showing the postures of 12h old larvae: control larvae (**E**) and upon activation of part of the TJ^+^ motoneuron population (**F**). **G** Number of peristaltic waves per 30 seconds at permissive (23°C) and restrictive (31°C) temperatures. Upon activation of 4 of the 6 TJ^+^ MNs, we observe a slight decrease in the number of peristaltic waves. Each single point represents a single 12h-old first instar larva. Error bars indicate the SD and n denotes the number of larvae tested. Statistical analysis: One way ANOVA. ***p≤0.001.

To confirm and further delineate the identity of TJ^+^ MNs we used *TJ-Gal4* and expressed a myristoylated targeted RFP reporter (*UAS-myr-RFP*) (generous gift from M. Landgraf). From late st16 to L3, *TJ-Gal4::UAS-myr-RFP* revealed the high reproducibility of these peripheral projections as well as their terminal processes onto muscles VO3-VO6 (supplementary Fig.3A-C) and onto the dorsal-most muscles LL1, DO5, DO2 and DO1 (from lateral to dorsal-most, supplementary Fig.3A’-C’). These results are in agreement with the above immunostaining results and we conclude that in each abdominal hemisegment there is a contingent of 2 TJ^+^/pMad^+^/*Islet-myc*^+^ MNs (ISNd MNs that project on muscles VO3-VO6), 3 TJ^+^/pMad^+^/Eve^+^ MNs (ISNdm MNs U1, U2 and U5, that respectively project on muscles DO1, D02 and LL1) and 1 TJ^+^/pMad^+^ MN that projects on muscle DO5.

To investigate the role of TJ^+^ MNs on locomotion we used *CQ2-LexA* (*13*) which allows specific activation of the three TJ^+^ U MNs U1, U2 and U5. Detailed monitoring of *CQ2-LexA* expression also revealed that this line additionally drives in mid-L1 stage in the TJ^+^ DO5-innervating MN (arrowhead in Fig.3D) and in 1 unidentified TJ^+^ IN located dorsally (not shown). Using *CQ2-LexA*, we therefore activated 4 of the 6 TJ^+^ MNs per hemisegment, and observed a moderate decrease in the number of peristaltic waves and no obvious locomotor phenotype nor body posture defect in the vast majority of the larvae (Fig.3G – second black bar). Importantly, no spastic paralysis was ever observed upon activation of TJ^+^ MNs using the *CQ2-LexA* driver (Fig.3E, F and video 4).

From this experiment we conclude that activation of more than 50% of TJ^+^ MNs does not trigger the spastic paralysis phenotype observed upon activation of the whole TJ^+^ population but rather causes a slight defect in locomotion probably due to the constant contraction of the dorsal muscles innervated by TJ^+^ MNs.

### TJ^+^-cholinergic and TJ^+^-GABAergic interneuron populations are distinctly involved in the locomotor behaviour of the larvae

Next we chose to use drivers allowing the targeting of specific interneuronal populations. We first investigated a possible expression of TJ in peptidergic, dopaminergic, serotonergic, and histaminergic INs and found no expression of TJ in these IN subclasses (supplementary Fig.4 C1-E2). Further molecular characterization using highly specific LexA drivers (*23*) showed that 10 out of 29 TJ^+^ neurons per hemisegment are cholinergic (Fig.4A-E’; supplementary Fig.5D-F1). Activation of those TJ^+^ cholinergic INs by intersectional genetics caused disrupted locomotor behaviour. Larvae frequently adopted a characteristic “crescent shape”, with head and tail regions brought close together by a tonic contraction of the ventral muscles (Fig.4T, video 5). This ventral contraction phenotype was heterogeneous in terms of severity, with some larvae being continually immobile with ventral contraction (video 5), while some others displayed bouts of ventral contraction interrupting otherwise seemingly normal crawling. Counting the number of peristaltic waves produced by these larvae reflected the heterogeneity of this phenotype, but also revealed that the number of waves were significantly reduced in these animals compared to control specimen (Fig.4F – second orange bar). Interestingly, such a spectrum of severity has also been recently reported when activation of different subsets of VNC neurons led to a ventral contraction phenotype (*9*).

**Fig. 4:**
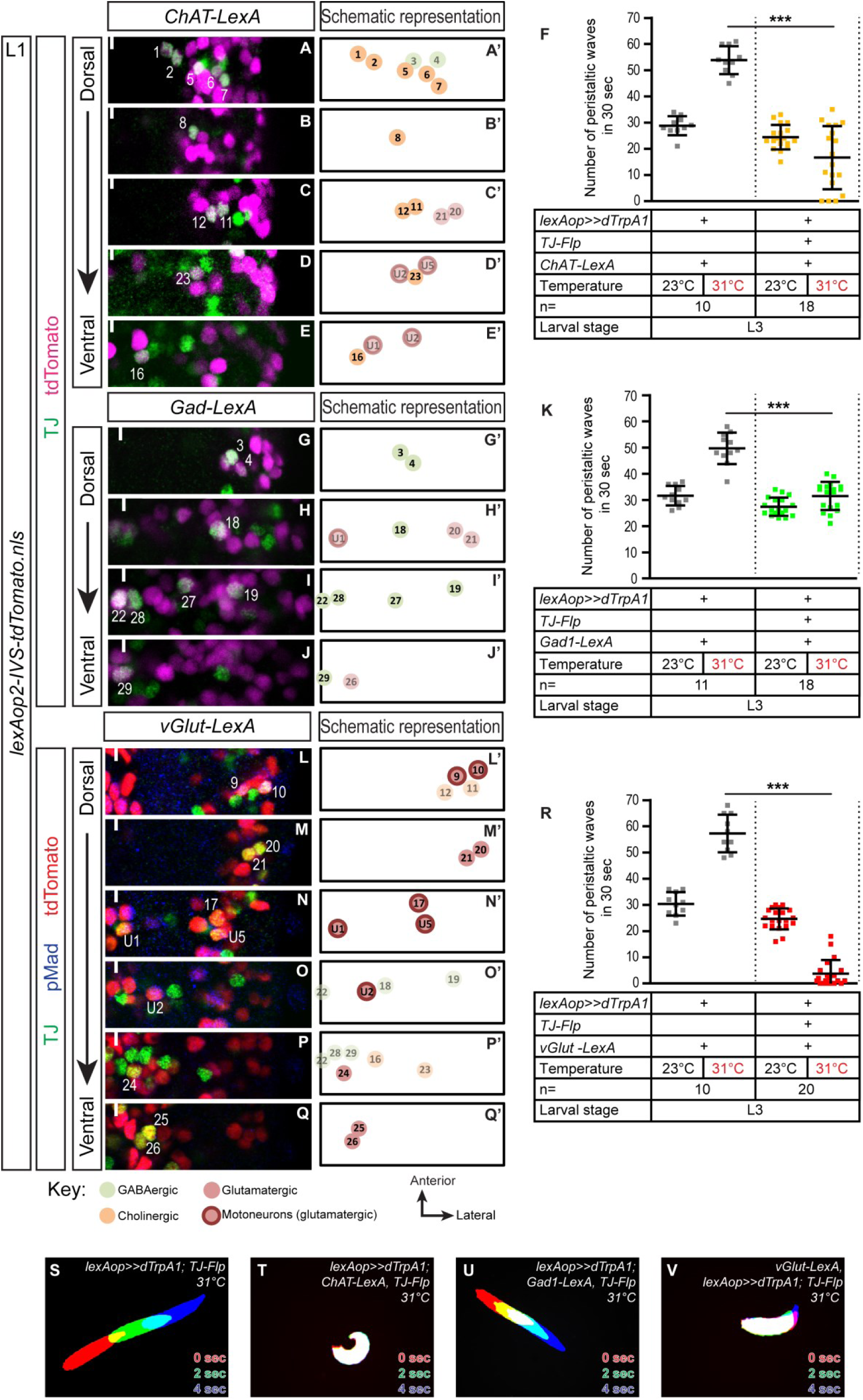
Distinct TJ^+^ subpopulations have distinct roles in the locomotor CPG and posture of larvae. **A-E, G-J, L-Q** Representative views of one hemisegment of a first instar larval VNC stained for TJ (green) and cholinergic cells (**A-E** - magenta – using *ChAT-LexA* driving *lexAop-nlsTomato*), or GABAergic cells (**G-J** - magenta – using *Gad1-LexA* driving *lexAop-nlsTomato*) or glutamatergic cells (**L-Q** - red – using *vGlut-LexA* driving a *lexAop-nlsTomato*) and the motoneuron marker pMad (blue). Cells are shown from dorsal (**A, G, L**) to ventral (**E, J, Q**) positions. The full line indicates the midline and anterior of the VNC is up. **A’-E’, G’-J’, L’-Q’** Schematic representation of the adjacent immunostaining panels. TJ^+^ cholinergic (**A’-E’**), GABAergic (**G’-J’**) and glutamatergic (**L’-Q’**) neurons are represented in bright colour while other non-cholinergic (**A’-E’**), non-GABAergic (**G’-J’**) and non-glutamatergic (**L’-Q’**) TJ^+^ neurons are shown with paler colours. The identity of those neurons is inferred from their position compared to TJ^+^ cholinergic, GABAergic or glutamatergic neurons. **F, K, R** Number of peristaltic waves per 30 seconds at permissive (23°C) and restrictive (31°C) temperatures of larvae expressing TrpA1 in TJ^+^ cholinergic neurons (**F**, orange bars), GABAergic neurons (**K**, orange bars) or glutamatergic (**R**, orange bars) neurons versus their respective controls that do not express TrpA1 (**F, K, R**, grey bars). Each single point represents a single third instar larva. Error bars indicate the SD and n the number of larvae tested. Statistical analysis: One way ANOVA. ***p≤0.001. **S-V** Superimposition of three consecutive time frames (0, 2 and 4 seconds) showing the postures of third instar larvae: control larvae (**S**) and upon activation of the TJ^+^ cholinergic (**T**), TJ^+^ GABAergic (**U**) and TJ^+^ glutamatergic (**V**) populations.

Among the remaining TJ^+^ INs, 8 per hemisegment are GABAergic (Fig.4G-J’; supplementary Fig.5A-C1). Examining *Gad1-LexA* co-localization with TJ, we noticed that among those 8 neurons, the 3 most ventral ones are located at the midline (neurons n°22, 28, 29 in Fig.4I-J’) and do not appear to have counterparts in the adjoining hemisegment, indicating that those cells are unpaired, midline cells (*24*). When activated, the TJ^+^-GABAergic IN subpopulation led to seemingly normal locomotion (Fig.4U, video 6). However, counting the number of peristaltic waves actually revealed that locomotion was slowed, with a reduction of 37% of the number of waves compared to control specimen at 31°C (Fig.4K - second green bar).

The remaining 5 TJ^+^ INs per hemisegment are glutamatergic and differentiating them from TJ^+^ MNs (also glutamatergic) is possible by counterstaining with pMad (Fig.4L-Q’; supplementary Fig.5G-I1). We chose to use *vGlut-LexA* in combination with *TJ-Flp* to activate this IN population, keeping in mind that by using such genetics we are also activating TJ^+^ MNs. We found upon activation of the entire TJ^+^ glutamatergic contingent a complete paralysis of the larvae, with forward propagating waves largely absent (Fig.4R – second red bar, Fig.4V). Another feature of this phenotype was the vertical lifting of the most anterior segments of the larvae (thoracic head region) off the substrate (video 7). As this phenotype is considerably different from the one observed upon activation of more than 50% of TJ^+^ MNs (see section above), we can assume that TJ^+^ glutamatergic INs play an important role in the normal locomotor behaviour of the larvae.

Taken together this data shows that activation of restricted subpopulations of TJ^+^ neurons defined on the basis of their neurotransmitter identity impact the crawling of the larvae with distinct behavioural hallmarks.

### A population of 3 TJ^+^ GABAergic INs per segment regulate the speed of locomotion

While searching for more restricted LexA drivers that would allow us to subdivide more finely the TJ^+^ population implicated in locomotion, we identified the *per-LexA* driver (*5*) whose expression co-localizes with 9 of the most ventral TJ^+^ neurons per segment (Fig.5A-F; neurons n°22,24,25,26,28 and 29 on Fig.1F and 1G). Activation of those 9 TJ^+^/*Per*^+^ neurons per segment led to a decrease in the number of peristaltic waves (Fig.5G – second purple bar, Fig.5I). Interestingly these larvae displayed relaxed, slightly atonic abdominal segments (video 8). Upon careful characterization of this TJ^+^/*Per*^+^ population, we discovered that 6 of them (3 per hemisegment) are glutamatergic (Fig.5D-F; neurons n°24, 25, 26 on Fig.1F and 1G) while the 3 remaining, located medially, are GABAergic (Fig.5A-C). Indeed, these three TJ^+^/*Per*^+^/GABAergic neurons correspond to the three unpaired, midline GABAergic cells that we described above (see Fig.4G-J; neurons n°22, 28, 29) and that do not have counterparts in the adjoining hemisegment. We were intrigued by those 3 unpaired neurons and used a sophisticated triple intersectional genetic approach based on the combinatorial expression of *TJ*, *Per* and *Gad1* to specifically activate them. A large proportion of the manipulated larvae (∼70%) displayed a marked reduction in their speed of locomotion compared to control larvae at 31°C (Fig.5J-L – video 9), while a minority appeared unaffected (∼30%). The incomplete or low dTrpA1 expression in every pool of 3 neurons in all segments for a given larva could explain the heterogeneity within the experimental group. In agreement with such possibility we noticed rather weak *Per-Gal4* expression in these neurons, which would be predicted to reduce the expression levels of the LexA^DBD^ component in this triple intersectional approach.

**Fig. 5:**
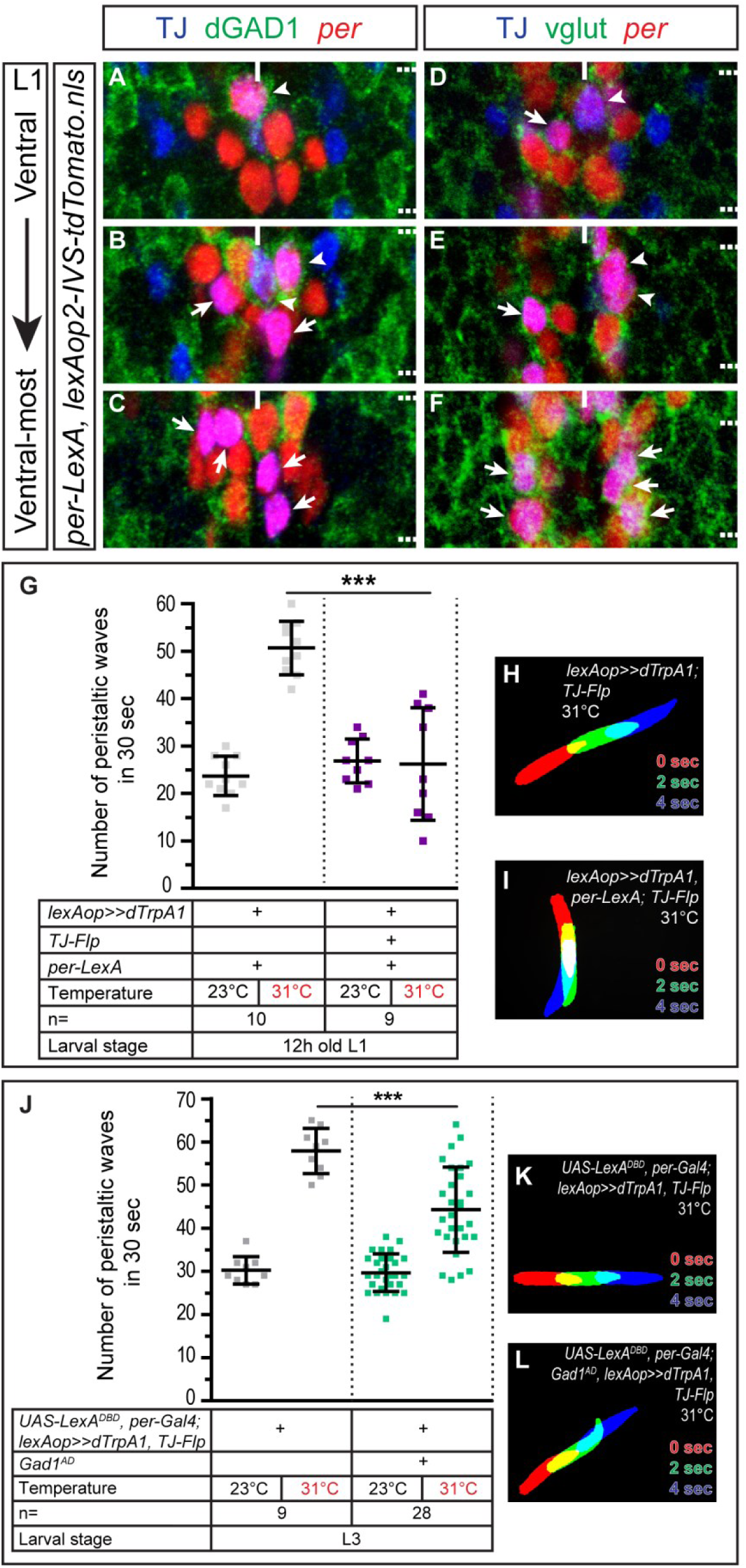
Three TJ^+^ *Per*^*+*^ GABAergic neurons per segment regulate the speed of locomotion in Drosophila larva. **A-C** Staining for TJ (blue), GAD1 (green) and nls-tdTomato (red) expressed with the *per-LexA* driver in freshly hatched first instar larva VNC. Staining reveals 9 TJ^+^ *Per*^+^ cells per segment, located in the ventral part of the VNC. The 3 most dorsal cells, located at the midline, are GABAergic (full arrowheads), while the 6 other cells are not (arrows). Note that one of the TJ^+^ *Per*^+^ GABAergic neurons (often the most dorsal) is weakly *Per*^+^. **D-F** Staining for TJ (blue), vglut (green) and nls-tdTomato (red) expressed under *per-LexA* driver in freshly hatched first instar VNC. The remaining 6 non-GABAergic cells are glutamatergic (arrows). **G** Number of peristaltic waves per 30 seconds at permissive (23°C) and restrictive (31°C) temperatures done by larvae expressing TrpA1 in TJ^+^ *Per*^+^ neurons (purple bars) versus controls that do not express TrpA1 (grey bars). Activation of the TJ^+^ *Per*^+^ population (9 cells per segments) lead to a drastic decrease in the speed of locomotion of 12h old larvae (second violet column). Phenotype is characterized by partial relaxed paralysis of the most-posterior segments of the larvae. One way ANOVA. ***p≤0.001. **H-I** Superimposition of three consecutive time frames (0, 2 and 4 seconds) showing the postures of third instar larvae: control larva (**H**) and upon activation of the TJ^+^ *Per*^*+*^ population (**I**). **J** Number of peristaltic waves per 30 seconds at permissive (23°C) and restrictive (31°C) temperatures done by larvae expressing TrpA1 in TJ^+^ *Per*^+^ GABAergic neurons (green-blue bars) versus controls that do not express TrpA1 (grey bars). Activation of the TJ^+^ *Per*^+^ GABAergic population (3 cells per segments) lead to a decrease in the speed of locomotion of third instar larvae (second green-blue column). One way ANOVA. ***p≤0.001. **K-L** Superimposition of three consecutive time frames (0, 2 and 4 seconds) showing the postures of third instar larvae: control larva (**K**) and upon activation of the TJ^+^ *Per*^*+*^ Gad1^+^ population (**L**).

Together these results show that activation of a population of 3 GABAergic TJ^+^/*Per*^*+*^ neurons per segment impacts the speed of locomotion in *Drosophila* larva.

### TJ^+^ *Per*^*+*^ GABAergic neurons, known as MNB progeny neurons, express a unique combination of TFs

Given the median position of the 3 GABAergic TJ^+^/*Per*^+^ neurons and the fact they lack counterparts in the adjoining hemisegment, we hypothesized that those neurons are a subset of the midline cells. Midline cells belong to the *sim* domain (*24*) and we confirmed by quadruple immunostaining with TJ, Gad1, Prospero and *sim-Gal4* driving a nuclear GFP that TJ^+^ median GABAergic neurons are *sim*^*+*^ in late embryonic stage 17 VNC (Fig.6A-B2, single and double empty arrowheads). We noticed weak *sim* expression in these neurons at this stage, an observation consistent with low *sim* expression in late embryonic stages as previously reported (*25*). It is important to note that the TJ^+^ non GABAergic (glutamatergic-positive) neurons located in the ventral part of the VNC are not *sim-Gal4*^*+*^ (depicted by arrows in Fig.6C-C2). We then found that 2 of the 3 TJ^+^ GABAergic midline cells belong to the Median Neuroblast (MNB) progeny subpopulation identified by nuclear Prospero expression (Prosp-nucl) (*24*) (Fig.6A, double empty arrowheads). We also found that all 3 TJ^+^ GABAergic cells are *fkh*^+^ and EN^+^ (Fig.6D-E, full arrowheads), two TFs known to be expressed in a subpopulation of MNB progeny but also iVUMs (*24*). To further delineate the exact identity of the third TJ^+^ GABAergic midline neuron (GAD^+^, *Per*^+^, *fkh*^+^, *En*^+^, Prosp^-^) we examined stage 16 embryo where midline cell identities can be determined accordingly to the highly stereotyped dorso-ventral and anterior-posterior location of the cells. Using these stereotyped positions along with *Per*, *fkh* and TJ immunostainings, we showed that TJ is not expressed in the iVUMs (supplementary Fig.6C and 6F - asterisks), easily recognizable by their ventral-most position among midline cells, located close to the posterior boundary of the segment and posterior to H-sib (supplementary Fig.6B and 6E - empty arrowhead). Instead we visualized TJ expression in cells located above the iVUMs in a position where the MNB progeny neurons are normally located (supplementary Fig.6B, C, E – circled cells and full arrowhead). We conclude that the 3 ventral TJ^+^ GABAergic INs are MNB progeny neurons and additionally show that those cells are singled out by their expression of the TFs *Hlh3b* (Fig.6G-I – full arrowheads) and *grain* (Fig.6J-M – full arrowheads). MNB progeny neurons arise from MNB neuroblast; strikingly, at stage 15 we found that this neuroblast expresses the TF *Jumu* (Fig.6N, double full arrowhead). Morphology of the MNB progeny neurons has only been scantily reported (*24, 26*). To visualize the anatomy of a single MNB progeny neuron per VNC, we took advantage of a *R47G08-lexA* driver we identified among the Janelia Farm Flylight collection and that co-localize with TJ specifically in 1 to 2 of the MNB progeny neurons per segment (data not shown). This lexA driver used in combination with *TJ-Flp* reported the characteristic morphology of single TJ^+^ MNB progeny neurons: from their cell body located ventrally they send a single neurite straight up dorsally, that then divides to follow both sides of the neuropile, spanning several segments anteriorly and posteriorly (Fig.6O, O’, O’’). To our knowledge this is the first description of MNB progeny neurons morphology at larval stages.

**Fig. 6:**
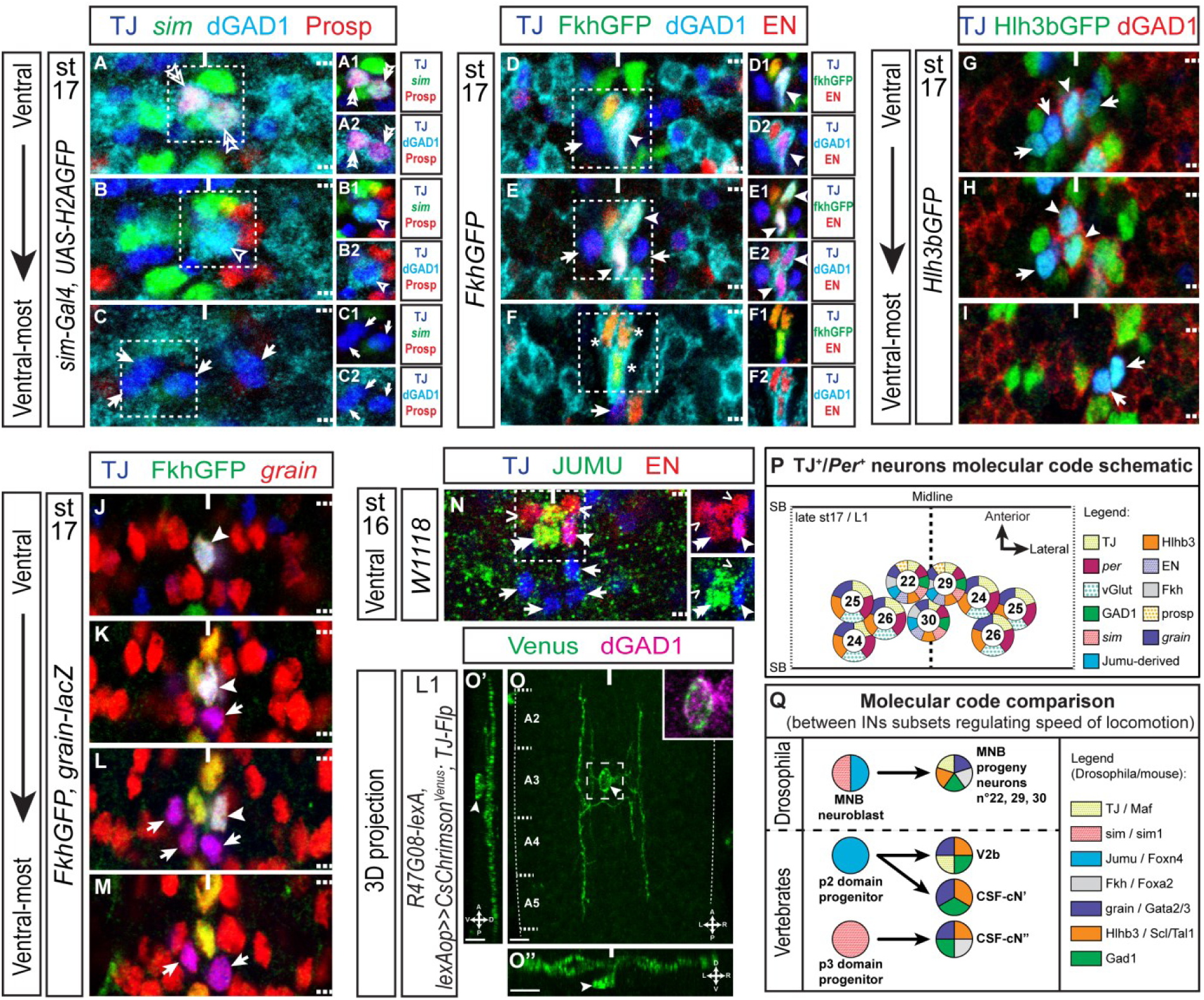
Molecular characterization of the 3 TJ^+^ *Per*^+^ GABAergic neurons identifies them as potential counterparts of the vertebrate CSF-cNs. **A-C** Staining for TJ (blue), GAD1 (cyan), Prospero (red) and GFP (green) driven by *sim-Gal4* in late stage 17 embryo VNC. *sim-Gal4* (green) is weakly expressed in GABAergic TJ^+^ ventral neurons only, identifying them as midline cells (**A** and **B,** simple and double empty arrowheads). Remaining TJ^+^ ventral population (TJ^+^ glutamatergic neurons – **C**, arrows – full contingent not shown) are *sim-Gal4*^-^, hence not part of the midline cells. Among the 3 TJ^+^ *sim*^*+weak*^ GABAergic located at the midline, two are positive for the MNB progeny neuron marker Prospero (**A**, double empty arrowheads) and one negative (**B**, simple empty arrowhead). **D-F** Staining for TJ (blue), GAD1 (cyan), Engrailed (red) and the fusion protein FkhGFP in stage 16 embryo VNC. TJ^+^ ventral GABAergic neurons (full arrowheads) are EN^+^ and Fkh^+^, two markers of MNB progeny subpopulation. Note in the ventral-most part of the VNC, located ventrally to TJ^+^ neurons, three EN^+^ Fkh^+^ Gad^+^ TJ^-^ cells: the iVUMs (asterisks in panel **F**). **G-I** Staining for TJ (blue), GAD1 (red) and Hlh3bGFP fusion protein (green) in late stage 17 embryo VNC. Both ventral GABAergic (full arrowhead) and glutamatergic (arrows) TJ^+^ neurons are Hlh3b^+^. **J-M** Staining for TJ (blue), FkhGFP fusion protein (green) and βgal (red) from *grain-lacZ* in late stage 17 embryo VNC. Both ventral GABAergic (full arrowhead) and glutamatergic (arrows) TJ^+^ neurons are *grain*^+^. **N** Staining for TJ (blue), JUMU (vert) and Engrailed (red) in stage 16 embryo VNC. The MNB neuroblast, identified by its position and size, is JUMU^+^ (double full arrowhead). EN^+^ TJ^-^ MNB progeny neurons (chevrons) and EN^+^ TJ^+^ MNB progeny neuron (full arrowhead) are closely affixed to the neuroblast. **O, O’, O’’** 3D reconstruction of a first instar larva VNC stained for dGAD1 (magenta) and showing the morphology of a single GABAergic TJ^+^ MNB progeny neuron (expressing Venus under the combined expression of *TJ-Flp* and *R47G08-lexA* driver, arrowhead) in dorsal view (**O**), lateral view (**O’**) and posterior view (**O’’**). For all three views (**O, O’, O”**), the Gad1 staining was removed to allow better observation of the MNB progeny neuron morphology. Close-up in O confirms that the TJ^+^ MNB progeny shown is indeed GABAergic. Scale bars: 5 µm. **P** Schematic representation of the molecular code for the most ventral *per*^*+*^ TJ^+^ neurons in late stage 17 embryo/L1 larva. A full segment is shown. **Q** Comparison of the molecular code found in TJ^+^ MNB progeny and the neuroblast they originate from and the CSF-cNs and the progenitors they arise from. TF code in vertebrates comes from (Andrzejczuk et al., 2018; Petracca et al., 2016).

Altogether we identify the TJ^+^ *Per*^*+*^ GABAergic population regulating the speed of locomotion as 3 MNB progeny neurons derived from a *Jumu*^*+*^ progenitor domain and singled out by the combinatorial expression of *TJ*, En, *fkh*, *Per*, *Hlh3b* and *grain.*

## Discussion

In this study, we characterized from embryogenesis to larval stage L3 TJ-expressing neurons in the VNC and investigated their role in the crawling behaviour of freely moving *Drosophila* larvae. We generated a *TJ-Flp* line and developed an intersectional genetic approach based on the use of TrpA1 to specifically activate different TJ^+^ subpopulations depending on their neurotransmitter properties.

### Charting the VNC: TJ, a reliable marker for a restricted subset of neurons in the embryo and larva

Our time course analysis revealed that TJ is expressed in the same restricted subset of neurons from the moment they are born in the embryo to the L3 larval stage. Our comprehensive mapping revealed that the number of TJ^+^ neurons (29 neurons/hemisegment) remains constant from st 17 to L3 within the abdominal A2-A6 region of the VNC. Within each hemisegment we counted 10 TJ^+^ cholinergic INs, 11 TJ^+^ glutamatergic neurons and 8 TJ^+^ GABAergic INs.

To our knowledge, such a detailed time course analysis of the expression profile of a TF within the VNC has been rarely described. One example is the highly restricted expression pattern of the B-H1/H2 homeoproteins in the VNC and their continuous expression within a small subpopulation of MNs identifiable from embryogenesis to late larval stages (*27*). In fact it is often a challenging task to precisely capture the complete spatial and temporal expression pattern of a gene of interest, particularly when it displays a broad expression pattern. Within the group of TFs expressed in restricted subpopulation of neurons in the VNC the bHLH gene *Dimmed* has been reported to be selectively expressed in central and peripheral neuroendocrine cells. Extensive work aimed at identifying and characterizing DIMM-expressing cells has generated a comprehensive map of nearly all 306 DIMM-positive cells in the central nervous system but the exact time course and minor changes of expression within these cells, although noticed, have not been systematically examined (*28*).

To date, one of the TFs that displays the most restricted and constant expression in the same group of VNC neurons is Eve (Even skipped). Eve is expressed from st13 onward in 16 cells per abdominal hemisegment: medially in the aCC, U1 and RP2 MNs and in the pCC IN; mediolaterally, in four U/CQ MNs (U2-U5); and laterally in eight to ten EL INs (*13, 29*). The identification of such cell type-specific TF and the characterisation of genetic tools such as Gal4 or LexA lines expression patterns of these “markers” have recently proved instrumental for characterizing neuronal circuits in the larval VNC. For example the recently elucidated neuronal circuit that promotes escape behaviour upon noxious stimuli in *Drosophila* larvae (*30*) involves the contribution of SNa MNs that were genetically amenable due to the highly specific *BarH1-gal4* line (*27*). Similarly, the very restricted *EL-Gal4* driver active in Eve-expressing lateral (EL) INs (*31*)capturing the has allowed deciphering the implication of EL INs within a sensorimotor circuit that maintains left-right symmetry of muscle contraction amplitude in the *Drosophila* larva (*13*). We thus foresee that our extensive mapping of TJ-expressing neurons, together with the reliable *TJ-Flp* line we generated will facilitate future studies aiming at identifying and investigating neuronal circuit formation and functioning in embryonic or larval VNC.

### TJ^+^ cholinergic neurons control body posture in *Drosophila* larva

Activation of the TJ^+^ cholinergic subpopulation gave rise to a ventral contraction phenotype, with larvae frequently adopting a “crescent shape” position. We noticed that the ventral contraction phenotype was heterogeneous between individuals. The larvae with the most dramatic features were persistently immobile and ventrally curved but peristaltic waves could still be observed emerging from the posterior part of the body (video 5). Another group of larvae displayed bouts of ventral contraction interrupting otherwise seemingly normal crawling phases that were characterized by regular propagation of peristaltic waves along the body. In light of these observations we currently favour the hypothesis that TJ^+^ cholinergic INs are part of a neuronal circuit controlling the body posture of the larva rather than being intrinsically involved in controlling the speed of locomotion.

Recently, Clark and collaborators (2016), while surveying Janelia Gal4 lines (*32*) crossed to *UAS-TrpA1*, identified from their screen several lines that gave rise to similar ventral contraction phenotypes. Interestingly, they also reported a spectrum of severity with some larvae continually blocked with tonically contracted ventral muscles, while others would go through periods of ventral contraction followed by attempts to crawl. From this screen three different lines specifically expressed in subsets of INs were identified and in the future it will be interesting to determine if these subsets include TJ^+^ cholinergic INs. Alternatively, it is possible that these INs subsets and TJ^+^ cholinergic INs are different components of the same circuit regulating ventral bending of the larva. In such a scenario we can foresee two major alternatives: 1) either these different IN subsets are linearly and sequentially activated, resulting in the contraction of the entire ventral muscle field via the activation of ISNb, ISNd and SNc MNs or 2) each IN subset is selectively and independently used as a premotor excitatory command allowing specific ventral groups of muscles to be activated via only one MN subpopulation (ISNb or ISNd or SNc). Such precise coordination of muscles contraction within a muscle field is exemplified by the recently described activity of an inhibitory IN denoted iIN. iIN specifically innervates “transverse” MNs and not “longitudinal” MNs, thus allowing for the sequential contraction of transverse and longitudinal muscles for an efficient contraction of the larval body segment (*15*).

### TJ^+^ glutamatergic neurons are components of a circuit controlling locomotion

Within each hemisegment, the TJ^+^ glutamatergic population comprises 6 MNs (U1, U2, U5, DO5 MN and 2 lateral Islet^+^ ISNd MNs), 3 ventral PMSI INs and 2 dorso-lateral INs. Our study on the functional role of TJ^+^ glutamatergic INs is hindered by the fact that when using *vGlut-LexA* both TJ^+^ MNs and TJ^+^ INs are targeted; to our knowledge no genetic tools exist that would allow us to specifically activate TJ^+^ glutamatergic INs. We nevertheless investigated the implication of TJ^+^ MNs using two non-optimal drivers (*CQ2-LexA and RapGAP1-LexA* (data not shown)) and found that TJ^+^ MNs activation leads to a defect in locomotion, i.e reduction of the number of peristaltic waves. Similarly, when we used *per-LexA* to activate the entire TJ^+^ PMSI population (6 TJ^+^ glutamatergic INs and 3 TJ^+^ GABAergic INs per segment) we noticed a reduction of the speed of locomotion and a partial relaxed paralysis of the posterior segments of the larvae. Interestingly, in both cases these genetic manipulations did not give rise to the severe spastic paralysis phenotype we observed while activating the entire TJ^+^ glutamatergic population. It could be argued that activation of the TJ^+^ PMSI GABAergic INs when using *per-LexA* is in some way “dominant” over the activation of TJ^+^ PMSI glutamatergic INs, and thus, it will be informative to only activate TJ^+^ PMSI glutamatergic INs. Unfortunately a *vGlut*^*AD*^ driver, an equivalent of the *GAD1*^*AD*^ line, has not been generated, thus precluding the implementation of this strategy at the present time. Nevertheless, it would be of interest to solely activate all *Per*^+^ glutamatergic INs (and thus not the *Per*^*+*^ TJ^+^ GABAergic INs); this could theoretically be achieved using the following transgenes: *lexAop>>dTrpA1*, *vGlut-LexA*, *per-Gal4* and an *UAS-Flp* (*33*). Since we reported strong expression of *per-Gal4* in the vast majority of *Per*^*+*^ glutamatergic neurons, we expect recombination events in the targeted neurons to be efficient and thus assume that this experiment will be conclusive.

### TJ^+^ GABAergic neurons regulate the speed of locomotion

Activation of TJ^+^ GABAergic neurons gave rise to an apparently normal, though slowed locomotion, with a number of peristaltic waves accomplished by larvae placed at 31°C similar to the number at 23°C. Since larvae tend to crawl faster when the temperature exceeds their 24°C comfort temperature (in young L3) (*34*), we wondered if TJ^+^ GABAergic neurons could be modulator component(s) of the temperature sensing system that detects uncomfortable temperatures and induces acceleration as a way to escape. TrpA1^+^ neurons located in the larval brain have been recently reported to be sensitive to the speed of the temperature increase (*35*) and we have found that these neurons do not express TJ (data not shown). The possibility that TJ^+^ GABAergic neurons are nonetheless part of this temperature sensing neuronal circuit awaits future experimental analysis.

Further subdivision within the TJ^+^ GABAergic INs pool, using a triple intersectional genetics approach, revealed that 3 *Per*^+^/TJ^+^ GABAergic INs located at the midline and known as MNB progeny neurons substantially impact the crawling speed of the larvae. It thus appears that this *Per*^*+*^ GABAergic population was overlooked in the previous characterization of PMSIs. This might be due to the fact that of the 20 *Per*^*+*^ INs present in each segment, only 3 are indeed GABAergic and these express low levels of *period*, especially in third instar larvae, a stage in which the characterization PMSIs was originally carried out (*5*).

### Molecular characterization of the *Per*^*+*^/TJ^+^ GABAergic (MNB progeny neurons); searching for equivalents throughout evolution

Cell body position and molecular characterization of the *Per*^+^ TJ^+^ GABAergic neurons allowed us to identify them as a subset of the MNB progeny among the midline cells. Although development of the midline cells has been meticulously described (*24-26*), the functional implication of these cells in locomotion or other behaviours is comparatively poor (*36, 37*). Here we show for the first time that MNB progeny neurons have a relevant function in the locomotor behaviour of the larva. Moreover, our studies add TJ as a new marker of the MNB progeny, offering the possibility to identify and genetically manipulate these neurons with *TJ-Gal4* or *TJ-Flp*.

Our detailed molecular characterization of the *Per*^+^ TJ^+^ GABAergic MNB progeny INs allowed us to survey for hypothetical counterparts in other model species. As reported here *Per*^+^ TJ^+^ GABAergic INs are characterized by their expression of *Hlh3b* and *grain*. In the mouse, the respective orthologous genes are *Tal1/Tal2/Scl* and *Gata2/3*, a transcription code specifically found in V_2b_ GABAergic INs (*16*). Interestingly, the V_2b_ IN subpopulation is known to regulate, in cooperation with V_1_ Ia INs, the limb CPG coordinating flexor-extensor motor activity (*38*). V_2b_ INs derive from a *Foxn4*^*+*^ progenitor domain (*39*); we find that the MNB neuroblast that gives rise to the MNB progeny neurons expresses *Jumu*, the *Drosophila* ortholog of *Foxn4*. Additionally, *Per*^+^ TJ^+^ GABAergic MNB progeny INs express *Fkh* and are midline cells derived from *sim*-expressing precursor cells. Intriguingly, a subset of V_2b_ INs in the mouse, also known as cerebrospinal fluid-contacting neurons (CSF-cNs, or KA, Kolmer-Agduhr neurons in Zebrafish) express *Foxa2*, the vertebrate orthologue gene of *Fkh* and arise from the *Sim1*^+^ progenitor domain p3 bordering the floor plate (see recapitulative schematic, Fig.6Q). This subset denoted CSF-cN″ is characterized by their very ventral location in the spinal cord abutting the floor plate (*40*). Since CSF-cN have been reported to modulate slow locomotion and body posture in Zebrafish (*3, 8*) it is tempting to speculate that *sim/jumu*-derived *Per*^+^/TJ^+^/*Fkh*^+^ MNB progeny neurons are the *Drosophila* counterpart of vertebrate *Sim1*-derived CSF-cN″ and *Foxn4*-derived CSF-cN’ neurons (See cartoon Fig.6Q). In the annelid *Platynereis dumerilii*, a recent study focusing on the molecular characterization of neuron types brought to light a group of neurons specifically co-expressing the TFs *Gata1/2/3* and *Tal* that may be related to CSF-cN (*41*) indicating that the molecular nature and physiological function of this neuronal type may have been conserved during evolution. The remarkable similarities of combinatorial expression of TFs within this IN class further exemplifies that the molecular mechanisms used during the wiring of the locomotor system are conserved and evolutionarily ancient.

## Materials and methods

### Antibody list

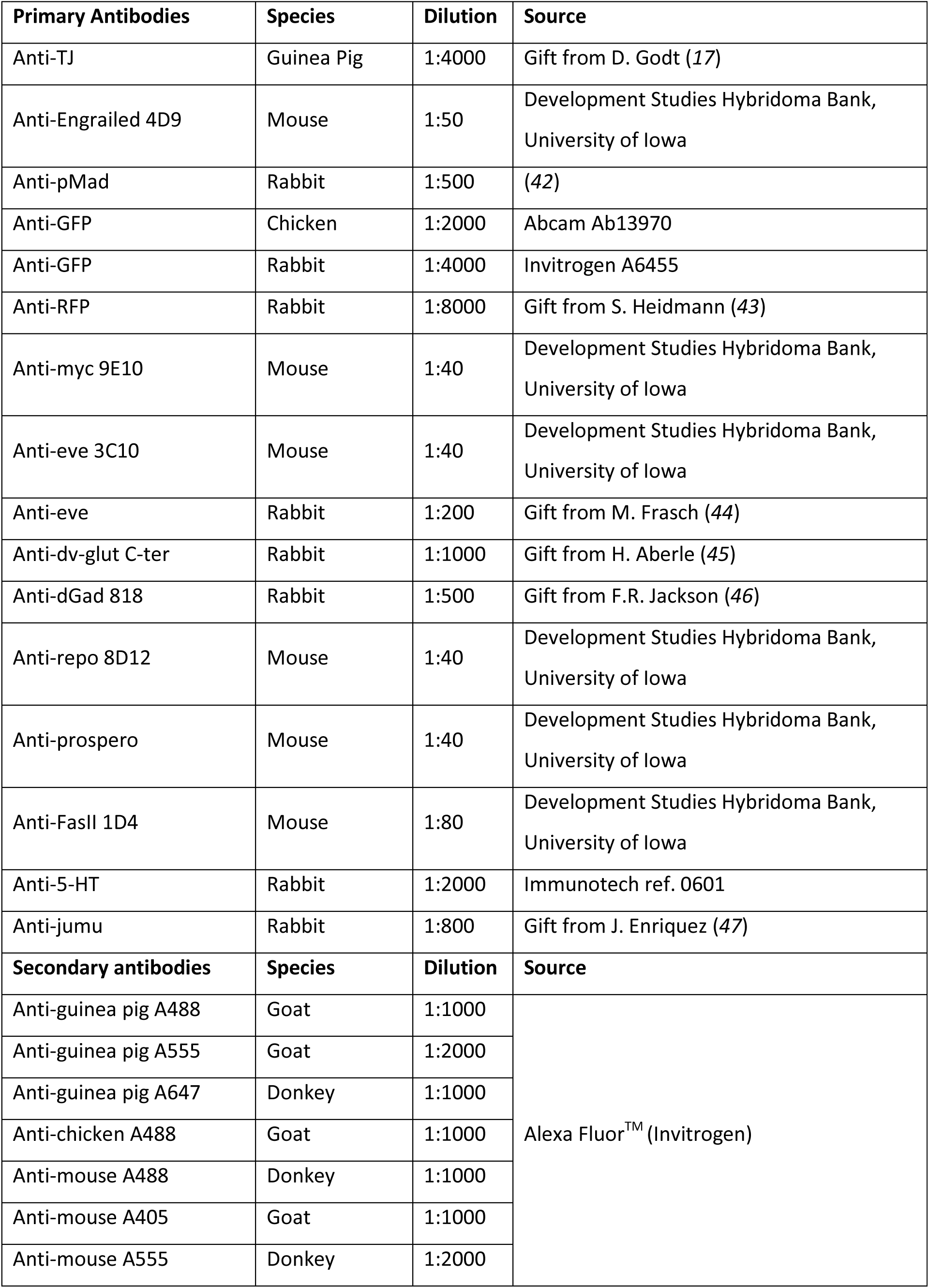
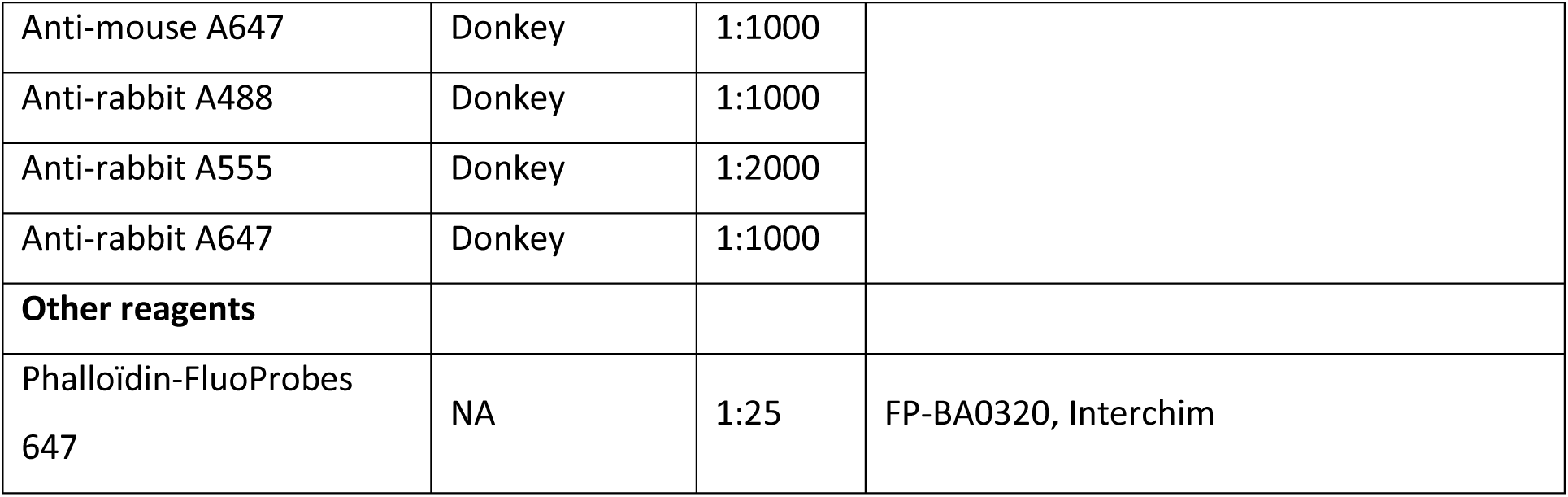

| Primary Antibodies | Species | Dilution | Source |
| --- | --- | --- | --- |
| Anti-TJ | Guinea Pig | 1:4000 | Gift from D. Godt (17) |
| Anti-Engrailed 4D9 | Mouse | 1:50 | Development Studies Hybridoma Bank,<br>University of Iowa |
| Anti-pMad | Rabbit | 1:500 | (42) |
| Anti-GFP | Chicken | 1:2000 | Abcam Ab13970 |
| Anti-GFP | Rabbit | 1:4000 | Invitrogen A6455 |
| Anti-RFP | Rabbit | 1:8000 | Gift from S. Heidmann (43) |
| Anti-myc 9E10 | Mouse | 1:40 | Development Studies Hybridoma Bank,<br>University of Iowa |
| Anti-eve 3C10 | Mouse | 1:40 | Development Studies Hybridoma Bank,<br>University of Iowa |
| Anti-eve | Rabbit | 1:200 | Gift from M. Frasch (44) |
| Anti-dv-glut C-ter | Rabbit | 1:1000 | Gift from H. Aberle (45) |
| Anti-dGad 818 | Rabbit | 1:500 | Gift from F.R. Jackson (46) |
| Anti-repo 8D12 | Mouse | 1:40 | Development Studies Hybridoma Bank,<br>University of Iowa |
| Anti-prospero | Mouse | 1:40 | Development Studies Hybridoma Bank,<br>University of Iowa |
| Anti-FasII 1D4 | Mouse | 1:80 | Development Studies Hybridoma Bank,<br>University of Iowa |
| Anti-5-HT | Rabbit | 1:2000 | Immunotech ref. 0601 |
| Anti-jumu | Rabbit | 1:800 | Gift from J. Enriquez (47) |
| Secondary antibodies | Species | Dilution | Source |
| Anti-guinea pig A488 | Goat | 1:1000 | Alexa Fluor™ (Invitrogen) |
| Anti-guinea pig A555 | Goat | 1:2000 |  |
| Anti-guinea pig A647 | Donkey | 1:1000 |  |
| Anti-chicken A488 | Goat | 1:1000 |  |
| Anti-mouse A488 | Donkey | 1:1000 |  |
| Anti-mouse A405 | Goat | 1:1000 |  |
| Anti-mouse A555 | Donkey | 1:2000 |  |
| Anti-mouse A647 | Donkey | 1:1000 |  |
| Anti-rabbit A488 | Donkey | 1:1000 |  |
| Anti-rabbit A555 | Donkey | 1:2000 |  |
| Anti-rabbit A647 | Donkey | 1:1000 |  |
| **Other reagents** |  |  |  |
| Phalloïdin-FluoProbes 647 | NA | 1:25 | FP-BA0320, Interchim |

### *Drosophila* stocks

| ***Drosophila* lines** | **Source** | **Identifier** |
| --- | --- | --- |
| *TJ-Gal4* | Tokyo Stock Center | DGRC # 104055 |
| *TJ-Flp* | generated by the lab | NA |
| *Gad1-LexA* | Bloomington Stock Center | BL # 60324 |
| *vGlut-LexA* | Bloomington Stock Center | BL # 60314 |
| *ChAT-LexA* | Bloomington Stock Center | BL # 60319 |
| *UAS-TrpA1* | Bloomington Stock Center | BL # 4308 |
| *UAS-shi<sup>ts</sup>* | (19) | NA |
| *lexAop>stop>dTrpA1* (2 lines on II<sup>d</sup> and III<sup>d</sup> chromosomes) | Obtained from the Rubin lab (Janelia Research Campus) | NA |
| *per-LexA* | Tokyo Stock Center | DGRC # 116999 |
| *per-Gal4* | Bloomington Stock Center | BL # 7127 |
| *FkhGFP* | Bloomington Stock Center | BL # 43951 |
| *Tsh-LexA* | Gift from J. Simpson (UC Santa Barbara)(48) | NA |
| *Tsh-Gal80* | Gift from G. Miesenböck (University of Oxford)(21) | NA |
| *lexAop-IVS-tdTomato.nls* | Bloomington Stock Center | BL # 66680 |
| *UAS-H2AGFP* | (49) | NA |
| *Act>>Gal4* | Bloomington Stock Center | BL # 4779 |
| *UAS-CD8GFP* | Bloomington Stock Center | BL # 5137 |
| *CQ2-LexA* | Gift from C. Doe (University of Oregon) (13) | NA |
| *sim-Gal4* (or *sim3.7-Gal4*) | Bloomington Stock Center | BL # 9150 |
| *Hlh3bGFP* | Gift from P. Tomancak (Max Planck Institute) (unpublished) | NA |
| *grain-lacZ* | (50) | NA |
| *Gad<sup>AD</sup>* | Bloomington Stock Center | BL # 60322 |
| *UAS-LexA<sup>DBD</sup>* | Bloomington Stock Center | BL # 56528 |
| *UAS-myrGFP* | Bloomington Stock Center | BL # 32198 |
| *B-H1-Gal4* | (51) | NA |
| *Tdc2-Gal4* | Bloomington Stock Center | BL # 9313 |
| *Dimm-Gal4* | Bloomington Stock Center | BL # 25373 |
| *TH-Gal4* (also called *ple-gal4*) | Bloomington Stock Center | BL # 8848 |
| *vmatGFP* | Bloomington Stock Center | BL # 60263 |
| *UAS>>TrpA1<sup>myc</sup>* | Bloomington Stock Center | BL # 66871 |
| *lexAop-Flp* | Bloomington Stock Center | BL # 55820 |
| *lexAop>stop>CsChrimson<sup>Venus</sup>* | Obtained from Rubin lab (Janelia Research Campus) | NA |
| *GMR47G08-LexA* | Bloomington Stock Center | BL # 52793 |

### Generation of the *TJ-Flippase*

#### Reagents

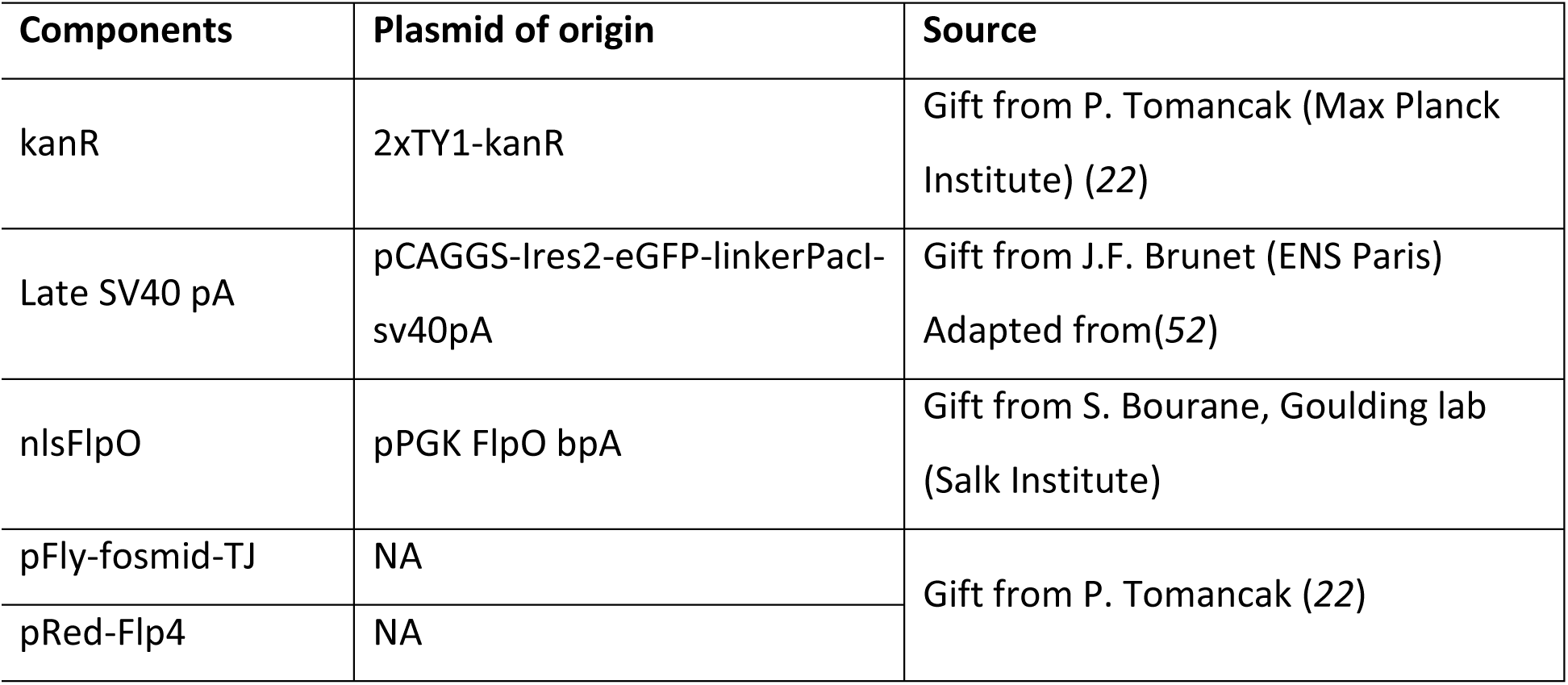
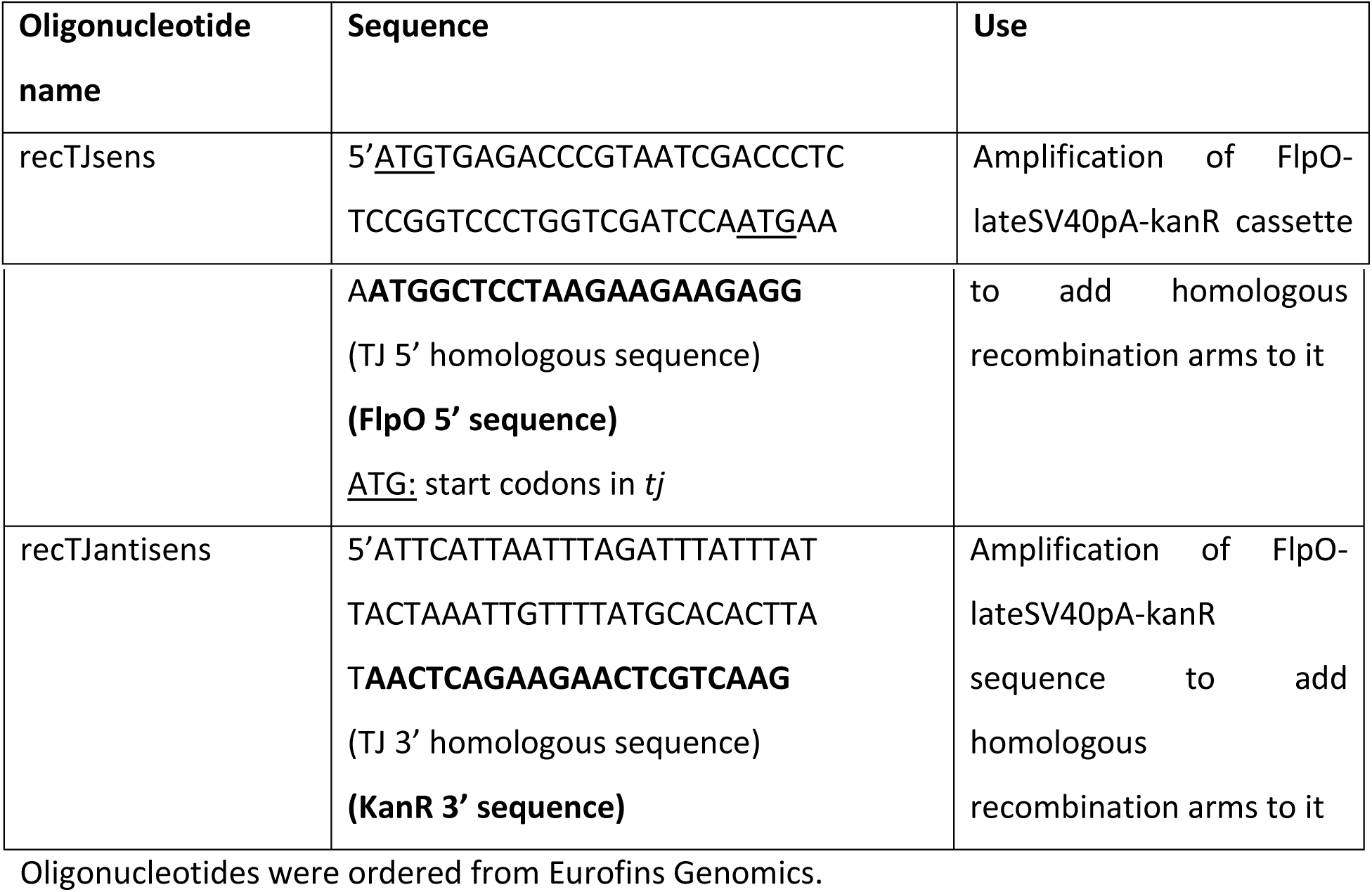

#### Protocol

We chose to use pFly-Fos technology (*22*) to generate *TJ-Flp* line after several trials at “transposon swap” strategies to replace gal4 transposon in *TJ-Gal4* line by a flippase failed. Ejsmont *et al* (2009) generated among their fosmid library a pFly-TJ-fosmid that contains the full *tj* sequence and probably all of *tj* regulatory elements; indeed expression of pFly-tj-Fos tagged with GFP perfectly recapitulates *tj* endogenous expression (data not shown). Briefly, in this fosmid and following Ejsmont and collaborators (2009) protocol, we replaced *tj* open reading frame and 3’ UTR by an optimized flippase sequence followed by a late SV40 polyadenylation sequence and a kanamycine resistance gene using the recombination oligonucleotides recTJsens and recTJantisens listed in the above table. Recombination oligonucleotides were chosen in order to: i) conserve the two endogenous *tj* transcription start signals (ATG) upstream of the flippase sequence in the *TJ-Flp* line and ii) conserve *tj* polyA signal sequence. Following recombination and contrary to Ejsmont protocol, we did not remove the kanamycine resistance sequence from the final fosmid used for transgenesis. Final fosmid was sent for targeted transgenesis in attp2 landing site (III^rd^ chromosome) (Bestgene Inc.).

### Immunohistochemistry

#### On larval VNC

Briefly, larvae of the right genotype were sorted out in Phosphate Buffered Saline-0,1% triton (DPBS 1X CaCl2^+^ MagCl2^+^ Gibco Invitrogen, Sigma-Aldrich; Triton X-100, Sigma Life Science) (PBS-T), rinsed in DPBS and transferred with tweezers to the DPBS-filled dissection chamber. Dissection chamber consists of a silicone-delineated well on poly-lysine coated glass slides; a double-sided piece of adhesive tape covers part of the dissection chamber floor. Dissection takes place on the adhesive tape. Head of the larvae were cut with scissors and posterior part of the body removed. Central Nervous Systems (CNS) were delicately dissected with tweezers (second and third instar larvae) or tungsten wires (first instar larvae) and all other tissues removed. CNS were then transferred to the non-adhesive covered part of the chamber and left to adhere to the slide, ventral part of the VNC against the slide (apart for period-observing dissections: CNS were placed dorsal part of the VNC against the slide). Further incubation steps were completed in the silicone chamber. CNS were fixed for 12 minutes with 3,7% formaldehyde diluted in DPBS (37% Formaldehyde solution, ref 252549, Sigma-Aldrich), then washed 3×5 minutes with PBS-T. At this point the double-sided piece of tape was removed. Blocking step was performed with PBS-T supplemented with 4% donkey serum (normal donkey serum S30-100 ml, Millipore) and 0.02% azide (Sodium azine NaN_3_, ref S2002, Sigma-Aldrich) for 30 minutes. CNS were then incubated with primary antibodies for 1h at room temperature (RT) or overnight at 4°C in a humid chamber, washed 3×5 minutes with PBS-T and then incubated with secondary antibodies 1h at RT in humid chamber in darkness. After 4×5 minutes of washing with PBS-T, silicone walls of the chamber were scraped off and Mowiol and a coverslip placed over stained CNS for imaging.

#### On larva VNC and muscle wall (for the study of MN projections)

For first instar larval (stL1) dissection embryos were preselected during the time when their main dorsal tracheae begin to fill with air, which represents 18 h after egg laying (AEL) and allowed to develop for further 3 h. First instar larvae (21 h AEL) were dissected as described in (*53*). Third instar larvae were heat-killed at 56°C for 5 sec and dissected as previously described in (*54*).

#### On embryo VNC

Immunolabeling of embryos was carried out as previously described (*55*).

### Image acquisition and processing

Images were acquired on a Zeiss LSM700 confocal with 40x or 63x objectives, treated and cropped in Photoshop (Adobe) and assembled in Illustrator (Adobe). For the benefit of colour-blind readers, double-labelled images were falsely coloured in Photoshop. 3D projection of whole VNC was implemented using Zen software (Zeiss).

### Locomotion assay

#### Larvae sorting

For first instar larvae testing, decorionated late stage 17 embryos of the right genotype were sorted out and placed on an agar/grape juice plate supplemented with some feeding medium (maize, sucrose and yeast). Hatching time was monitored and larvae locomotion assessed 6 hours after hatching.

For third instar larvae testing, eggs were laid for 5 hours on basic maize feeding medium. Approximately 72 hours later burrowing third instar larvae were picked up and assessed for locomotion.

#### Assay

Larvae clean of food were gently picked up with tweezers (in the case of the third instar larvae) or the back of tweezers (for first instar larvae) and placed on a 56 mm-agar plate supplemented with grape juice. After a 30-seconds acclimation period, the number of peristaltic waves done by the larvae was manually counted (at that time plate surface temperature was 23°C). The plate was transferred on a hot plate to heat for 2 minutes and 30 seconds or until it reached 31°C. The plate was quickly removed from the heat and the number of peristaltic waves done by the larvae manually assessed for 30 seconds more. Plate was left to rest for 4 minutes until surface temperature reached 23°C. Number of peristaltic waves in 30 sec was assessed once more.

### Statistical analysis

Statistical tests were carried out using Graphpad Prism (Graphpad software, Inc). We used one-way ANOVA with a Tukey post-hoc test to analyse more than two groups of data. When comparing only two groups we used unpaired Student t-test.

## Acknowledgments

We thank the Developmental Studies Hybridoma Bank at the University of Iowa, the Bloomington Stock Center and the *Drosophila* Genetic Resource Center in Kyoto Institute of Technology for monoclonal antibodies and fly stocks. We would also like to thank D. Godt, S. Heidmann, M. Frasch, H. Aberle, F.R. Jackson, M. Landgraf, Y. Aso and G. Rubin, J. Simpson, G. Miesenböck, C. Doe, P. Tomancak, S. Bourane, J. Enriquez and S. Baulac for generously sharing fly lines, antibodies and vectors. This work was funded by grants from INSERM and a 3-year PhD funding from the Association Française contre les Myopathies (AFM) for H.B (Doctoral funding n°19408). Imaging analysis was carried out on the regional reference core facility (RIO) supported by the French Ministry of Scientific Research.

## Author contributions

A.G., P.C. and H.B. designed the research. H.B. carried out experiments and processed the data presented in all figures but Fig.3A-C, Supp Fig.1, Supp Fig.3 and Supp Fig.4. A.G. carried out the experiments leading to the results presented in Fig.3A-C, Supp Fig.1, Supp Fig.3 and Supp Fig.4. C.S. produced preliminary data for the project under the supervision of S.Y. and J.B.T. J.V. financially supported the beginning of the project. A.G. and H.B. wrote the manuscript. P.C., J.B.T. and C.S reviewed the manuscript.

## Additional information

The authors declare no competing interests.

